# Use of Droplet Digital PCR for Consistent Detection of TMPRSS2:ERG Gene Fusion Transcripts Initiated In Vitro

**DOI:** 10.1101/2025.03.26.645503

**Authors:** Megan H. Check, Sarah E. Ernst, Karen S. Sfanos

**Affiliations:** Department of Pathology, Johns Hopkins University School of Medicine, Baltimore, MD, USA; Sidney Kimmel Comprehensive Cancer Center, Baltimore, MD; Department of Urology, Johns Hopkins University School of Medicine, Baltimore, MD

**Keywords:** Prostate cancer, gene fusions, TMPRSS2, ERG, TNFα, inflammation

## Abstract

Gene fusions are hybrid genes that arise from chromosomal rearrangements linking two independent genes. The most common gene fusion in prostate cancer involves the 5’ androgen-regulated TMPRSS2 promoter fused with the 3’ ETS transcription factor ERG. TMPRSS2:ERG (T:E) gene fusions occur in about half of all prostate cancers and are considered an early event in oncogenesis. Investigations into the mechanism behind T:E gene fusion initiation using *in vitro* systems are hindered by the technical limitations posed by fluorescence *in situ* hybridization and suboptimal sensitivity of reverse transcription quantitative PCR. The objective of this study was to develop a reliable, user-friendly method of detecting low abundance T:E gene fusion transcripts as a read-out for putative T:E gene fusions generated in cells. We identified droplet digital PCR (ddPCR) as a sensitive method for detecting rare T:E gene fusion transcripts and observed consistent detection in reactions containing a single T:E gene fragment, or 1 fusion positive cell per 10,000 fusion negative cells. Next, we evaluated dihydrotestosterone (DHT), genotoxic insults (irradiation, etoposide), and inflammatory agents tumor necrosis factor-alpha (TNFα) and hydrogen peroxide (H_2_O_2_) as initiators of T:E fusions in a fusion negative prostate cell line (LNCaP). Consistent with prior studies, we identified DHT combined with etoposide as potent, synergistic initiators of T:E gene fusion transcript expression in LNCaP cells. We did not detect T:E gene fusion transcripts after LNCaP exposure to irradiation, TNFα, or H_2_O_2_. We determined that TNFα and H_2_O_2_ exposure led to global downregulation of androgen receptor signaling, which may have limited the formation or expression of treatment-initiated genomic T:E fusions. Therefore, one limitation of the ddPCR assay is the requirement for T:E fusion mRNA expression. Our proposed method significantly improves the feasibility of testing novel initiators of the T:E gene fusion and can be applied to additional studies investigating mechanisms of gene fusion initiation in prostate and other cancers.

## Introduction

Gene fusions occur through chromosomal rearrangements linking two independent genes into one hybrid gene that produces a hybrid mRNA transcript. One key example of gene fusions that frequently occur in prostate cancer are those that involve 5’ genomic regulatory elements controlled by androgen signaling fused to members of the ETS-domain transcription factors.^1^ The most common gene fusion in prostate cancer involves the 5’ untranslated region of the transmembrane serine protease 2 (TMPRSS2) gene joined to the protein-coding portion of ETS transcription factor ERG (ERG), a growth-promoting ETS-transcription factor.^2^ Here, the transcription of ERG becomes androgen regulated, leading to constitutive expression of ERG in prostate cancer cells that promotes cellular proliferation^3^ and a stem-cell like phenotype.^4^ The TMPRSS2:ERG (T:E) gene fusion is the most common chromosomal rearrangement in prostate cancer that occurs in up to half of all prostate cancers, although its prevalence can differ by race or ethnic group.^5–8^ The T:E gene fusion is thought to occur early in prostate carcinogenesis due to the homogeneity of ERG expression in tumor foci, reflecting clonal expansion of a parent cell containing the gene fusion.^9^ T:E fusions have likewise been identified, albeit rarely, in putative prostate cancer precursor lesions such as high grade prostatic intraepithelial neoplasia (HGPIN) and proliferative inflammatory atrophy (PIA), providing additional evidence for the early carcinogenic role of T:E fusions in prostate epithelial cells.^10,11^

Several factors are hypothesized to contribute to the generation of T:E gene fusions in prostate cells, including androgen receptor signaling, double-stranded DNA (dsDNA) breaks, and inflammatory mediators. Testosterone and dihydrotestosterone (DHT) activate androgen signaling in cells that express the androgen receptor (AR) protein. Active signaling through AR drives the relaxation of chromatin around AR-regulated genes, such as TMPRSS2, and also induces chromosomal proximity of the TMPRSS2 and ERG loci.^12,13^ Initiation of the gene fusion depends on dsDNA breaks occurring at both the TMPRSS2 promoter and the ERG gene. One proposed mechanism for dsDNA break generation that contributes to T:E fusion formation is through topoisomerase II beta (TOP2B). TOP2B is an enzyme that relieves torsional strain by the cleavage and religation of dsDNA during active transcription. In fact, TOP2B is an essential component of AR-mediated transcription and is recruited with AR to androgen-regulated promoters.^14^ T:E breakpoints were found to be in close proximity to TOP2B cleavage sites initiated during active AR signaling, providing additional evidence for the role of TOP2B in T:E fusion formation.^14^ Poisons of TOP2B (i.e., etoposide) and ionizing radiation are other sources of dsDNA breaks leading to T:E fusions^12–14^, though these results stem from *in vitro* studies, and these factors likely do not contribute to fusions initiated in the human prostate. Recently, prostate inflammation has been investigated for contributing to the initiation of T:E gene fusions. One report showed that the exposure of prostate cancer cell lines to tumor necrosis factor alpha (TNFα) *in vitro* generated T:E gene fusions through a free radical-dependent mechanism.^15^ We likewise identified rare instances of T:E gene fusions occurring in the luminal epithelial cells of the inflammation-associated lesion PIA in the setting of bacterial prostatitis.^10^ Cumulatively, this evidence suggests a potential role for inflammation in driving the generation of the T:E gene fusion as an early oncogenic event.

Research into the initiating factors of T:E gene fusions using fusion-negative prostate cells is hampered by the fact that T:E fusions are initiated at a very low rate *in vitro*, occurring at a frequency of 1 per 10,000 cells.^13,14^ Detecting infrequent T:E fusions in a population of fusion-negative prostate cells is technically challenging. Two techniques to detect putative T:E gene fusions *in vitro* have been previously used: (1) fluorescence *in situ* hybridization (FISH) and (2) reverse transcription quantitative PCR (RT-qPCR). For FISH, fluorophore-tagged DNA probes hybridize to complementary sequences within TMPRSS2 loci, ERG loci, and the intervening region, which allows the visualization of chromosome 21 and its integrity at these sites. Chromosomal rearrangements or dsDNA breaks at these sites can be visualized by adjacent fluorescent signals “breaking-apart” or disappearing as that sequence of DNA is lost. Though many studies have used this technique to suggest fusion initiation *in vitro*, it only shows that chromosomal rearrangements occurred and does not definitively prove that a viable T:E gene fusion took place. Likewise, the visualization of rare T:E fusions (presumably 1 per 10,000 cells) is technically difficult using FISH, especially given that the interpretation of FISH imaging can be influenced by how each cell has been longitudinally sectioned, and ideally requires confirmation of similar patterns in neighboring cells. Additionally, FISH is time intensive and requires trained personnel for interpretation.

RT-qPCR detects the expression of T:E fusion transcripts at the RNA level, which is often used as a proxy for putative genomic T:E gene fusions. Transcripts are typically amplified with primers designed against exon 1 of TMPRSS2 and exon 4 of ERG, the most common exon-exon junction in T:E gene fusion transcripts.^12,16^ Several studies have used RT-qPCR^13–15,17^ to measure T:E fusion transcripts following exposure to DHT, irradiation, or etoposide in fusion negative prostate cancer cell lines (LNCaP, LAPC-4). A major drawback of RT-qPCR for detecting the expression of T:E gene fusions is its low sensitivity. Strategies to enhance detection include enriching for fusion transcripts in starting material and screening a large amount of sample material. Fusion transcripts can be enriched by using ERG-specific primers during RNA reverse transcription.^14^ Another approach is nested RT-qPCR, which involves two subsequent rounds of amplification for T:E fusion transcripts.^17^ Other groups have measured fusion transcript expression by screening a substantial amount of biological replicates (∼20-50) and by dividing samples into nearly 200 individual RT-qPCR reactions.^13–15^ These approaches are time consuming, labor intensive, and invite error or misinterpretation as multiple rounds of PCR are performed.

The current study aimed to develop an alternative method for measuring T:E gene fusions initiated *in vitro* that is highly sensitive and user-friendly. Droplet digital PCR (ddPCR) was selected as a platform based on its reliability and validity in detecting low abundance targets. The development of a reliable and reproducible method for the detection and measurement of experimentally-induced gene fusions will enable future studies aimed at assessing potential exogenous initiators or modulators of T:E gene fusion expression *in vitro*.

## Materials and Methods

### Cell culture

LNCaP cells (Cat. No. CRL-1740, American Tissue Culture Collection [ATCC]) were cultured in RPMI medium with 10% fetal bovine serum (FBS; Gibco), VCaP cells (Cat. No. CRL-2876, ATCC) were cultured in DMEM medium with 10% FBS. Mycoplasma testing and short-tandem repeat cell line testing were performed at regular intervals. LNCaP cells were grown to 50-70% confluence in T25 flasks prior to use in experiments.

### RNA isolation and cDNA synthesis

RNA was isolated from cultured cells using TRIzol Reagent (Cat. No. 15596026, Invitrogen) and the manufacturer’s instructions with the following modification: an additional wash with 75% ethanol was performed at the RNA wash step. RNA was quantified using the Qubit RNA Broad Range kit (Cat. No. Q32856, ThermoFisher). cDNA was synthesized from 5 ug of RNA using oligo(dT) primers and the SuperScript III First-Strand Synthesis System (Cat. No. 18080051, ThermoFisher). cDNA samples intended for ddPCR underwent clean-up using the DNA Clean & Concentrator-25 kit (Cat. No. D4034, Zymo Research).

### ddPCR

Reactions were assembled using ddPCR supermix (Cat. No. 1863024, Bio-Rad), cDNA, and TaqMan assays for the *T:E fusion* (FAM, Assay ID: Hs03063375_ft) and *TBP* (VIC, Assay ID: Hs00427620_m1). Each reaction was divided into eight to eleven technical replicates (20 uL/well). Every run included a negative control (LNCaP cells, steady state), a positive control (LNCaP cells exposed to DHT and etoposide), and a no template control (NTC). Droplets were generated using the QX200 Droplet Generator (Bio-Rad) according to the manufacturer’s instructions. Droplets were transferred into a 96 well plate (Cat. No. 12001925, Bio-Rad) and PCR was subsequently performed on the Applied Biosystems Veriti Thermal Cycler (Thermo Fisher Scientific). The PCR cycle conditions can be found in **Table S1**. The plate containing the droplets was transferred to the QX200 Droplet Reader (Bio-Rad) which assessed droplets for fluorescent signal.

### ddPCR analysis

Data was analyzed using QX Manager Software Standard Edition (2.1.0). Technical replicate wells were combined for analysis using the merge-well function in the QX Manager software. Wells containing less than 10,000 droplets or low-quality droplet clusters were excluded from analysis. A positive signal for T:E gene fusion detection required that ≥3 FAM positive droplets were detected. Thresholds for positive droplets in FAM and VIC channels were calibrated through droplet amplitudes in positive control wells using the built-in algorithm within QX Manager Software. Statistical analysis was performed using GraphPad Prism 10.

### RT-qPCR

RT-qPCR was performed using iTaq Universal Probes Supermix (Cat. No. 1725132, Bio-Rad), diluted cDNA (1:20 in DNA-free water), and TaqMan assays for *AR* (Assay ID: Hs00171172_m1), *NF-κB* (Assay ID: Hs00765730_m1), *NKX3*.*1* (Assay ID: Hs00171834_m1), *TMPRSS2* (Assay ID: Hs01122322_m1), and *TBP* (Assay ID: Hs00427620_m1). Each reaction was performed with three technical replicates. Fluorescence intensity was measured using the CFX Connect Real-Time PCR Detection System (Bio-Rad). The PCR cycle conditions can be found in **Table S2**. Relative gene expression was calculated via the ΔΔCt (2^-ΔΔCt^) method relative to the reference gene TATA-box binding protein (*TBP*).

### Standard curve using T:E fusion gene fragment

T:E gene fusion DNA fragments (gBlocks, IDT, **Table S3**) were quantified using the Qubit DNA High Sensitivity kit (Cat. No. Q32854, ThermoFisher) and serially diluted in DNA-free water containing poly(A) (Cat. No. 10108626001, Roche) to ensure fragment stability. One microliter of serially diluted T:E gene fragments (1-20 copies/µL) was added to ddPCR reactions containing 2 µL cDNA (approximately 200 ng/µL) from LNCaP cells. Poly(A) solution was used as a negative control. Reactions were divided into eight technical replicates and ddPCR was performed as described above.

### Standard curve using VCaP cells spiked into LNCaP cells

LNCaP and VCaP cells were counted using the Countess Automated Cell Counter (ThermoFisher). VCaP cells (20-200,000 cells) were spiked into 2 million LNCaP cells to achieve varying ratios (1:10-1:100,000) of VCaP to LNCaP cells. RNA was extracted as described above.

### Androgen deprivation followed by exposure to DHT, etoposide, or irradiation

LNCaP cells were grown in RPMI media containing 10% charcoal-stripped FBS (CSS; Corning) for 48 hours prior to the addition of DHT (Cat. No. D-073, Sigma-Aldrich), methanol vehicle (Cat. No. 34860, Sigma-Aldrich), or etoposide (Cat. No. S1225, Selleck Chemical). Irradiation was performed using the CIXD XStrahl irradiator.

### TNFα and hydrogen peroxide (H_2_O_2_) exposure

TNFα (Cat. No. H8916, Sigma-Aldrich) was reconstituted in molecular biology-grade water, diluted into cell culture media containing 10% FBS, and added to LNCaP cells for 48 hours. Hydrogen peroxide (Cat. No. H1009, Sigma-Aldrich) was diluted in PBS before being added to the cell layer.

## Results

### Evaluation of the ddPCR assay for detecting T:E gene fusion transcripts at low abundance

As an initial step, we aimed to evaluate the sensitivity of ddPCR in detecting T:E gene fusion transcripts. Our assay uses TaqMan primers and probe that are specific to fusions occurring between exon 1 of TMPRSS2 and exon 4 of ERG. To test the sensitivity of the assay, we serially diluted DNA fragments containing the T:E gene fusion (**Table S3**) to concentrations between 1 and 20 copies per µL, then added 1 µL of each dilution to ddPCR reactions (20 µL) containing cDNA from the T:E fusion-negative LNCaP prostate cancer cell line. We consistently detected T:E gene fragments as low as 1 copy per 20 µL reaction (0.05 ± 0.01 copies/µL) in all biological replicates (**Table 1, Fig. 1A**). Our findings suggest the ddPCR assay reliably detects T:E gene fragments at very low abundance, with detection capability approaching a single T:E copy per reaction.

**Table 1.** Detection of *TMPRSS2:ERG* (T:E) Gene Fragments by Droplet Digital PCR.

| **T:E gene fragment copies per 20 uL reaction** | **Expected concentration (copies/uL)** | **Measured concentration (copies/uL ± SD)** |
| --- | --- | --- |
| 20 copies | 1.00 | 0.70 ± 0.11 |
| 10 copies | 0.50 | 0.36 ± 0.03 |
| 5 copies | 0.25 | 0.14 ± 0.04 |
| 2 copies | 0.10 | 0.08 ± 0.00 |
| 1 copy | 0.05 | 0.05 ± 0.01 |
| 0 copies | 0.00 | 0.00 ± 0.00 |
| NTC | 0.00 | 0.00 ± 0.00 |

*TMPRSS2:ERG* (T:E) gene fragments were diluted to low abundance and detected using Droplet Digital PCR (ddPCR). The expected concentration was determined by dividing the theoretical number of input copies by total reaction volume (20 uL). The measured concentration was determined through ddPCR. The average and standard deviation of three biological replicates is reported. NTC = no template control. SD = standard deviation.

**Figure 1.**
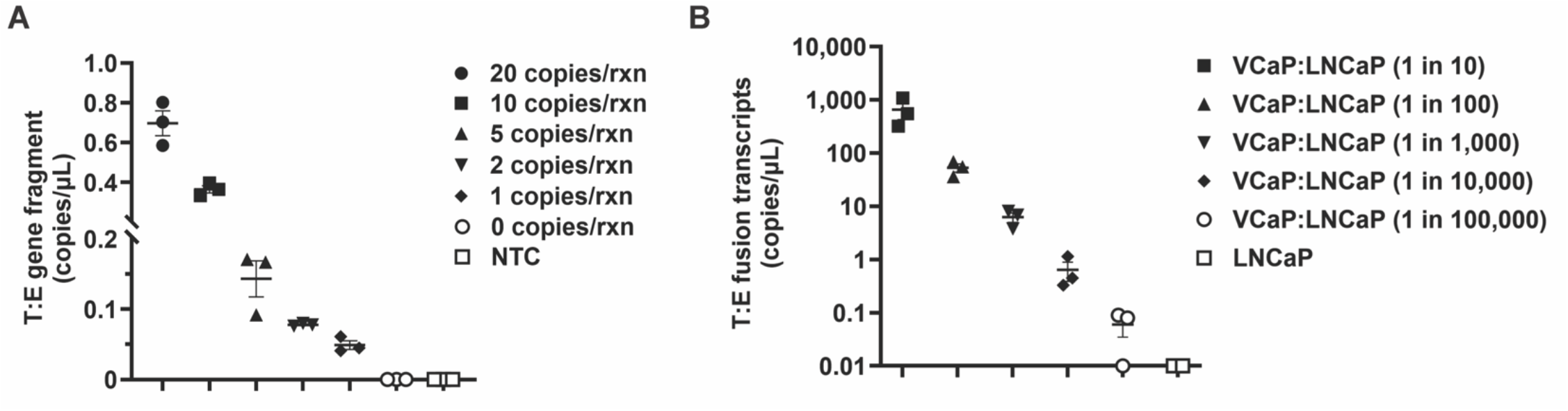
ddPCR consistently detects TMPRSS2:ERG gene fusions at low abundance. **A**) DNA fragments containing the TMPRSS2:ERG (T:E) gene fusion were serially diluted to low abundance (1-20 copies/uL) and added to ddPCR reactions containing cDNA from T:E fusion-negative LNCaP cells. The legend displays the expected number of input T:E gene fragment copies per 20 uL reaction. The concentration of T:E gene fragments was determined using ddPCR. Data is representative of 3 biological replicates. Concentration is displayed as mean +/-SEM. **B**) T:E gene fusion transcripts are consistently detected by ddPCR in cell populations containing a frequency of down to 0.01% (1 in 10,000) fusion-positive cells (VCaP) in a background of fusion-negative cells (LNCaP). The concentration of T:E fusion transcripts was determined using ddPCR. Data is representative of 3 biological replicates. Concentration is displayed as mean +/-SEM on a log-scale. Data containing a concentration of zero (0 copies/uL) is displayed as 0.01 copies/uL here to enable graphing on a log-scale.

As a second approach to assessing the sensitivity of the ddPCR assay, we sought to determine the sensitivity of ddPCR for detecting T:E gene fusion transcripts expressed by a T:E fusion-positive cell in the background of T:E fusion-negative cells. For this experiment, we spiked a prostate cancer cell line positive for the T:E gene fusion (VCaP) into prostate cancer cells negative for the gene fusion (LNCaP). Varying numbers of VCaP cells were mixed with a uniform amount of LNCaP cells to achieve different ratios of VCaP to LNCaP cells (1:10 – 1:100,000). Fusion transcripts were consistently detected in all VCaP spiked samples from ratios of 1:10-1:10,000 (VCaP:LNCaP cells, **Fig. 1B**). This result demonstrated that ddPCR reliably detects fusion transcripts at their estimated rate of initiation *in vitro* (1 in 10,000 cells).^13,14^ Additionally, we were able to detect fusion transcripts in samples containing a ratio of 1:100,000 (VCaP:LNCaP cells, **Fig. 1B**). This result occurred in 2 out of 3 biological replicates, which demonstrates that 1 fusion-positive cell per 100,000 fusion-negative cells approached the limit of sensitivity of our ddPCR assay.

### Consistent generation of T:E gene fusions following exposure to etoposide and DHT

Next, we sought to test the ability of the ddPCR assay to recapitulate previously published data for the initiation of T:E gene fusions in LNCaP cells *in vitro* using DHT, ionizing radiation, and etoposide.^13^ Prior to the experimental exposures, LNCaP cells underwent androgen deprivation for 48 hours in media containing charcoal stripped serum (CSS), according to previously reported methods for T:E fusion intiation.^13,14^ Simulating an environment deprived of androgens created a baseline to observe the effect of DHT towards the initiation of T:E gene fusions in an experimental setting. LNCaP cells were simultaneously exposed to DHT (100 nM) or methanol (vehicle) and a genotoxic insult in the form of ionizing radiation or etoposide, a potent TOP2B poison. The cells were incubated for 48 hours prior to RNA harvest. Fusion transcripts were not detected in LNCaP cells exposed to DHT or vehicle alone (UT, **Fig. 2**). In contrast, T:E fusion transcripts were consistently detected across biological replicates in LNCaP cells exposed to DHT with etoposide (**Fig. 2**). The transcript copy number likewise increased as the dose of etoposide increased from 50 µM to 100 µM. Surprisingly, no fusion transcripts were detected in LNCaP cells exposed to ionizing radiation (5 Gy or 10 Gy) with or without DHT present (**Fig. 2**).

**Figure 2.**
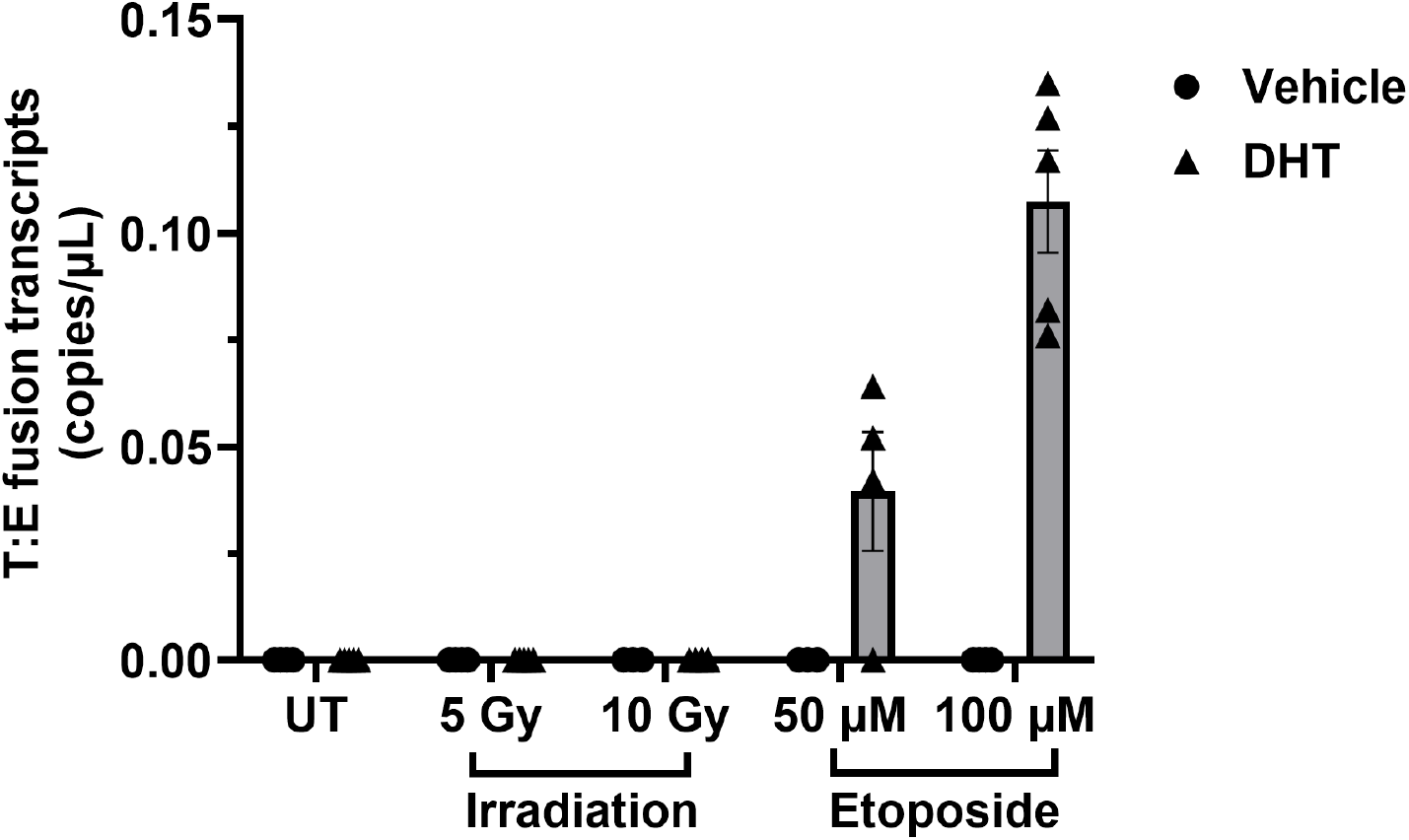
TMPRSS2:ERG gene fusions are consistently generated in a fusion-negative cell line following exposure to dihydrotestosterone (DHT) combined with etoposide. LNCaP cells (fusion-negative) underwent 48 hours of androgen deprivation with RPMI + 10% charcoal-stripped serum, followed by 48 hours of treatment with DHT (100 nM) or methanol (vehicle) combined with ionizing radiation (5 or 10 Gy), or etoposide (50 or 100 µM). Exposure to DHT combined with etoposide consistently generated T:E gene fusion transcripts, as detected by ddPCR. ddPCR reactions were divided into eleven technical replicates to screen the cDNA sample in its entirety (92% of 5 ug). Data is representative of 3-5 biological replicates. Concentration is displayed as mean +/-SEM. UT = untreated control that did not receive irradiation or etoposide.

### TNFα exposure does not lead to detectable T:E fusion expression and decreases AR signaling

We next tested the effects of inflammatory stimuli on T:E fusion gene expression in LNCaP cells. First, we tested TNFα exposure as an inciter of fusion gene expression based on a previous report.^15^ LNCaP cells were exposed to TNFα for 48 hours prior to RNA harvest. No fusion transcripts were detected after TNFα exposure alone at 10 ng/mL and 100 ng/mL concentrations (**Fig. 3A**). Likewise, the addition of DHT (100 nM) during TNFα treatment (100 ng/mL) did not facilitate expression of T:E gene fusions, as it did in the positive control of DHT + etoposide (**Fig. 3A**).

**Figure 3.**
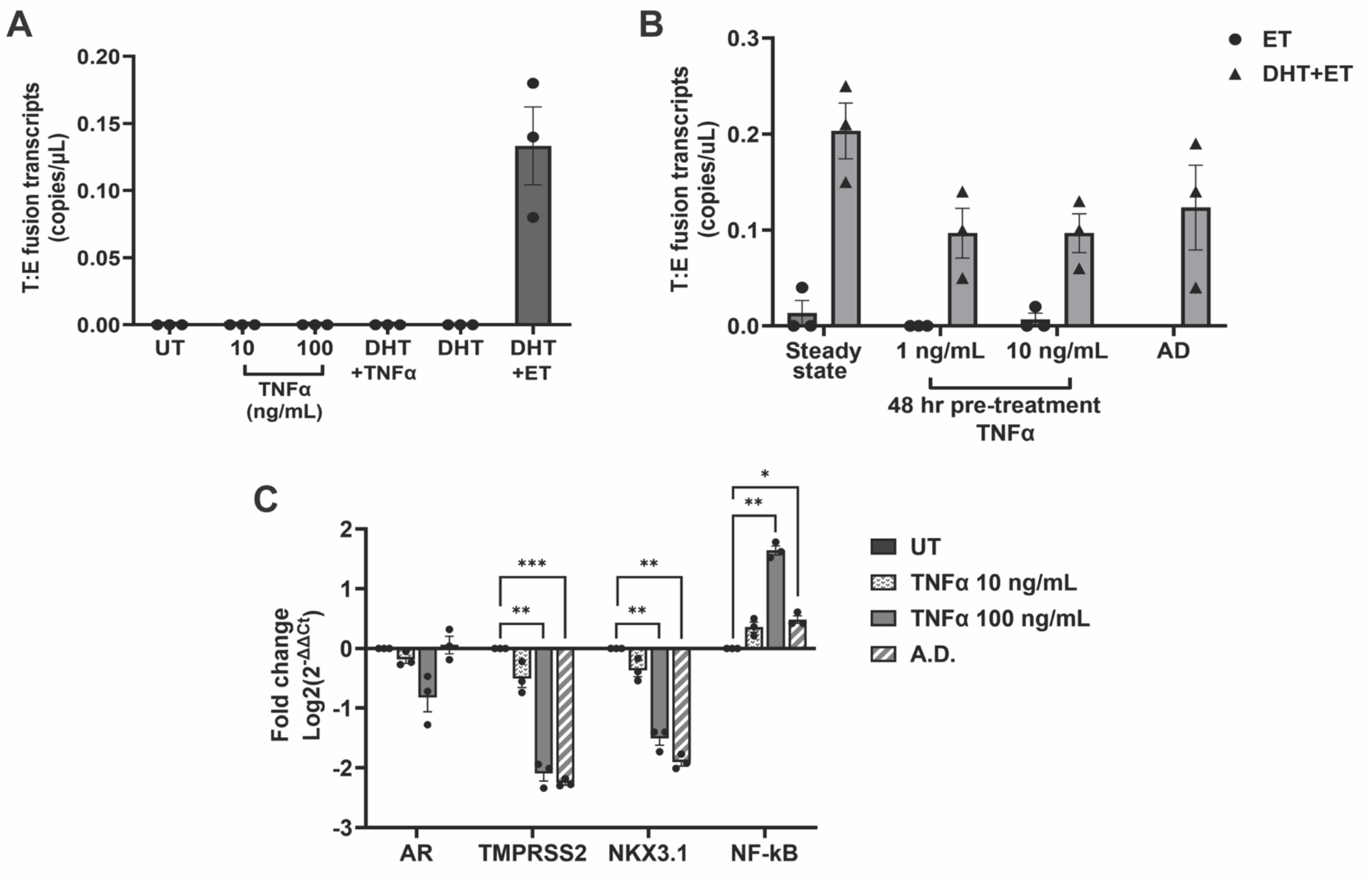
TNFα exposure does not lead to detectable T:E fusion expression in LNCaP cells and decreases androgen receptor (AR) expression. **A**) T:E gene fusion transcripts were not detected after LNCaP cells were exposed to TNFα (10 or 100 ng/mL) alone or with a combination of TNFα (100 ng/mL) and DHT (100 nM) for 48 hours. LNCaP exposure to DHT (100 nM) and etoposide (100 µM) for 48 hours served as a positive control (DHT+ET) for T:E gene fusion generation. The concentration of T:E gene fusion transcripts was determined using ddPCR. ddPCR reactions were divided into eleven technical replicates to screen the cDNA sample in its entirety (92% of 5 ug). Data is representative of 3 biological replicates. Concentration is displayed as mean +/-SEM. **B**) T:E gene fusion transcript levels generated by DHT and etoposide (DHT+ET) are lower in LNCaP cells pre-treated with TNFα. LNCaP cells were pre-treated with TNFα (1 or 10 ng/mL), androgen deprivation (AD, RPMI + 10% CSS), or steady state media (RPMI + 10% FBS) for 48 hours prior to exposure to DHT (100 nM) and etoposide (100 µM) for T:E gene fusion generation. The concentration of T:E gene fusion transcripts was determined by ddPCR. ddPCR reactions were divided into eleven technical replicates to screen the cDNA sample in its entirety (92% of 5 ug). Data is representative of 3 biological replicates. Concentration is displayed as mean +/-SEM. **C**) TNFα reduces expression of androgen receptor (*AR*) and androgen-regulated genes (*TMPRSS2, NKX3*.*1*) in LNCaP cells. LNCaP cells were exposed to TNFα (10 or 100 ng/mL) or androgen deprivation (AD, RPMI + 10% CSS) media for 48 hours prior to RNA harvest. Gene expression was measured via qRT-PCR using Taqman assays (*AR, TMPRSS2, NKX3*.*1, NF-κB, TBP*). *NF-κB* was used as a positive control. Log2 fold change is relative to reference gene, *TBP*. Data is representative of 3 biological replicates. Data are displayed as mean +/-SEM. Statistical significance was assessed using a one sample t test, with significance set at *p* < 0.05. (\**p* < 0.05, \*\**p* < 0.01, \*\*\**p* < 0.001). AD = androgen deprivation, UT = untreated control.

Next, we questioned whether the expression of the T:E gene fusions elicited by DHT and etoposide was affected by pre-exposure to TNFα. We hypothesized that TNFα may induce reactive oxygen species within LNCaP cells that may facilitate fusion generation by DHT and etoposide. For this experiment, we exposed LNCaP cells to TNFα (1 ng/mL and 10 ng/mL, respectively) for 48 hours prior to exposure to DHT and etoposide for 48 hours. We also included a group that was exposed to androgen deprivation for 48 hours prior to DHT and etoposide exposure. Interestingly, the expression of gene fusion transcripts was decreased in all samples pretreated with TNFα, to a similar degree as androgen deprivation conditions (**Fig. 3B**). When normalized to the housekeeping gene (TBP), pre-treatment with TNFα (10 ng/mL) induced significantly lower T:E fusion transcripts than DHT + etoposide without TNFα pre-treatment (**Fig. S1**).

Due to this finding, we questioned whether treatment of LNCaP cells with TNFα leads to lowered AR signaling, as a possible explanation for a decrease in fusion transcript expression. We performed RT-qPCR for AR and the AR-regulated genes TMPRSS2 and NK3 homeobox 1 (NKX3.1) as well as the downstream activation target of TNFα, nuclear factor kappa B subunit 1 (NF-κB), on LNCaP cells that were exposed to TNFα (10 or 100 ng/mL) for 48 hours. We found that treatment with TNFα led to a marked decrease in *AR, TMPRSS2*, and *NKX3*.*1* expression levels, and, as expected, an increase in *NF-κB* (**Fig. 3C**).

### Hydrogen peroxide exposure does not lead to detectable T:E fusion gene expression

Finally, we sought to test the effects of free radicals on the generation of T:E gene fusions. A previous study reported that the fusion gene-inciting effects of TNFα were dependent on reactive oxygen species^15^, therefore we aimed to test this by exposing LNCaP cells to H_2_O_2_. H_2_O_2_ is a common experimental source of free radicals owing to its release of hydroxyl radicals (HO^-^) during its decomposition into water and oxygen. After determining the highest tolerable doses of H_2_O_2_ on LNCaP cell viability (Fig. S2), we exposed LNCaP cells to H_2_O_2_ (100 µM or 500 µM), with and without DHT, and allowed the cells to incubate for 48 hours. No fusion transcripts were detected for all LNCaP cells treated with H_2_O_2_, despite the detection of fusion transcripts in the positive control combination of DHT + etoposide (**Fig. 4A**). We likewise demonstrate that exposure of LNCaP cells to H_2_O_2_ (500 µM) led to a decrease in *AR* expression (**Fig. 4B**).

**Figure 4.**
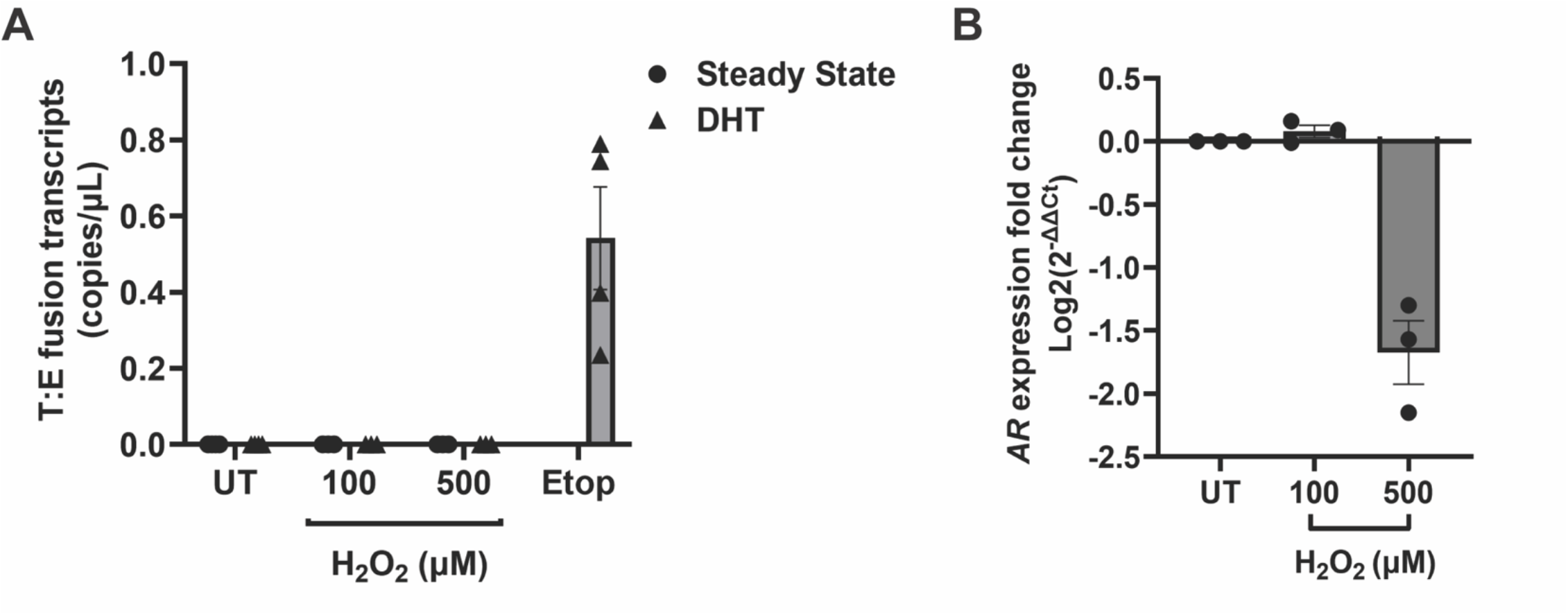
T:E gene fusion transcripts were not detected after LNCaP cells were exposed to H_2_O_2_. **A**) LNCaP cells were exposed to H_2_O_2_ (100 or 500 µM) for 48 hours with and without the addition of DHT (100 µM) to steady state media (RPMI + 10% FBS). LNCaP exposure to DHT (100 nM) and etoposide (100 µM) for 48 hours served as a positive control for T:E gene fusion generation. The concentration of T:E gene fusion transcripts was determined using ddPCR. ddPCR reactions were divided into eleven technical replicates to screen the cDNA sample in its entirety (92% of 5 ug). Data is representative of 3-4 biological replicates. Concentration is displayed as mean +/-SEM. Etop = etoposide. **B**) H_2_O_2_ reduces expression of the androgen receptor (AR) gene in LNCaP cells. LNCaP cells were exposed to H_2_O_2_ (100 or 500 µM) for 24 hours prior to RNA harvest. Gene expression was detected through qRT-PCR using Taqman assays (*AR, TBP*). Log2 fold change is relative to reference gene, *TBP*. Data is representative of 3 biological replicates. Concentration is displayed as mean +/-SEM. UT = untreated control.

## Discussion

The objective of this study was to develop a reliable, user-friendly method for detecting low abundance T:E gene fusion transcript expression as a read-out for T:E fusion gene generation in cells after genotoxic exposures. The results of this study demonstrate that ddPCR consistently detects T:E gene fusions at an estimated one copy per reaction in the background of cDNA from fusion-negative cells. Additionally, ddPCR reliably quantified T:E gene fusion transcripts expressed by cell populations containing 1 fusion-positive cell per 10,000 fusion-negative cells. Consistent with the literature, this study also found that DHT and etoposide were potent, synergistic initiators of T:E gene fusion transcript expression in LNCaP cells.^13,14^ This finding was consistent across many biological replicates, and therefore continued to serve as the positive control for our assay. In contrast to previous findings, however, T:E fusion transcripts were not detected after fusion-negative LNCaP cells were exposed to irradiation, TNFα, or H_2_O_2_.

Our finding that etoposide exposure required concurrent administration of DHT to generate T:E fusions (**Fig. 2**) provides further evidence that activation of AR signaling in the presence of a DNA damaging agent leads to the generation of T:E gene fusions. DHT-mediated activation of AR signaling promotes the co-localization of TMPRSS2 and ERG loci, bringing them into spatial proximity.^12,13^ Expression of the AR protein is essential for this co-localization to occur, as cell lines that are negative for AR (AR-), such as DU145, do not show co-localization of TMPRSS2 and ERG loci following stimulation with DHT.^17^ Nevertheless, the results of our study suggest that short term DHT exposure (48 hours) alone was not sufficient to induce T:E gene fusions in LNCaP cells (**Fig. 2**).

AR-mediated signaling recruits TOP2B to active sites of transcription at AR-regulated genes, including TMPRSS2.^14^ TOP2B relieves torsional strain by creating a dsDNA break that is quickly followed by re-ligation of the DNA ends. TOP2B is essential to the expression of AR-regulated genes and constitutes at least one source of naturally-occurring dsDNA breaks at the TMPRSS2 loci.^14^ Etoposide, a chemical poison of TOP2B, generated abundant fusion transcripts when it was combined with DHT exposure in our assay (**Fig. 2**). Etoposide exposure alone during androgen starved conditions (without the addition of DHT) did not lead to detectable T:E gene fusion transcripts in LNCaP cells (**Fig. 2**). This finding is not conclusive, however, as T:E gene fusions may have been formed at the DNA level, but not expressed due to global suppression of AR signaling by androgen deprivation. It is of interest to note, however, that etoposide treatment of LNCaP cells without androgen deprivation (**Fig. 3B**, steady state) also did not lead to consistent expression of T:E fusion transcripts.

We also found that irradiation did not result in the detection of T:E gene fusion transcripts in LNCaP cells (**Fig. 2**). A previous study identified fusion gene transcripts after irradiating LNCaP cells (50 Gy) and exposing them to DHT-containing media (100 nM) for 24 hours.^13^ In our study, we irradiated the LNCaP cells at lower doses (5 Gy and 10 Gy), which may have been insufficient to effectively initiate T:E gene fusions and lead to the expression of T:E gene fusion transcripts. Another key difference is that we allowed the cells to incubate for 48 hours, instead of 24, after irradiation and DHT exposure. Even so, it is unlikely that a longer incubation time reduced the detection of fusion transcripts, as we consistently detected fusion transcripts in other samples that underwent 48 hour incubation.

An unexpected finding from this study was that TNFα or H_2_O_2_ exposure did not lead to expression of T:E gene fusion transcripts in the fusion-negative LNCaP cell line (**Fig. 3A,. 4A**). This finding is contrary to a previous study that demonstrated that TNFα as well as TNFα-induced free radicals initiated T:E gene fusion transcripts in fusion-negative cells.^15^ Detection of T:E gene fusion transcripts is dependent on the expression level of the TMPRSS2 gene promoter. We determined that TNFα significantly reduced the expression of AR-regulated genes, including *TMPRSS2* and *NKX3*.*1*, in the LNCaP cell line (**Fig. 3C**). This result is consistent with previous reports.^18,19^ In the context of the present study, TNFα-mediated downregulation of TMPRSS2 likely prevented the expression, and thus detection, of putative T:E gene fusion transcripts. We combined DHT with TNFα treatment to determine whether we could upregulate any existing TNFα-generated gene fusions, but still found no evidence for T:E gene fusion transcripts (**Fig. 3A**). Additionally, our data shows that pre-treatment with TNFα reduced the expression of T:E gene fusion transcripts induced by DHT + etoposide, to a similar degree as androgen deprivation pre-treatment (**Fig. 3B**). We speculate that TNFα may deter the initiation of T:E gene fusions *in vitro* due to the global downregulation of AR-mediated signaling, an essential component of T:E gene fusion initiation. We identified a similar degree of dampened AR-mediated signaling when LNCaP cells were exposed to H_2_O_2_ (500 µM), indicating a similar explanation for our lack of detection of T:E gene fusion transcripts with H_2_O_2_ exposure (**Fig. 4A,B**).

We also found that androgen deprivation is not required prior to DHT + etoposide for the induction of T:E gene fusions in LNCaP cells. Previous studies demonstrating the initiation of T:E gene fusions *in vitro* implemented androgen deprivation for two to three days prior to exposure to DHT.^13,14^ Here, we observed that more T:E gene fusion transcripts were elicited after DHT + etoposide exposure when cells were maintained in steady state media compared to androgen deprivation media (**Fig. 3B**), though this trend did not reach statistical significance (**Fig. S1**). Accordingly, we ceased pre-treatment with androgen deprivation media in subsequent experiments and observed a greater amount of T:E fusion transcripts expressed after the positive control treatment (DHT + etoposide, **Fig. 4A**).

A key limitation of this study is that our method of gene fusion detection depends on the expression of T:E gene fusion mRNA transcripts. Whereas our method detects the generation of authentic gene fusions at the mRNA level, it is unable to assess fusions that have occurred at the DNA level and that are not expressed due to, for example, lowered AR signaling. Another limitation is that the expression of T:E gene fusion transcripts may not translate into protein-coding fusion transcripts, as the protein-coding exons in ERG can sometimes be lost in the chromosomal rearrangements leading to T:E gene fusions.^20^ Techniques used to detect T:E gene fusions at the DNA or protein level require that a large amount of material be screened to obtain appreciable levels. Assaying for T:E gene fusions at the RNA transcript level theoretically allows the greatest chance of detecting gene fusions generated *in vitro*. A single T:E gene fusion on one allele may be transcribed many times, potentially generating multiple T:E fusion transcripts that can be detected through ddPCR. However, DNA and protein techniques can be employed with increased odds of success once optimal stimuli are determined for T:E gene fusion transcript expression.

## Conclusions

We demonstrate that ddPCR is a sensitive, user-friendly method for detecting rare T:E gene fusions generated *in vitro*. We confirmed that etoposide and DHT are synergistic promoters of T:E gene fusions and found that irradiation (at 5 Gy and 10 Gy), TNFα, or H_2_O_2_ did not generate detectable T:E gene fusions. Inflammatory mediators (TNFα, H_2_O_2_) should be further investigated for having a potential role in T:E fusion generation, but in the context of globally suppressed AR signaling. Our method significantly improves the feasibility of testing novel initiators of the T:E gene fusion and can be applied to studies investigating the mechanism of other gene fusions in prostate cancer.

## Acknowledgements

This work was supported by the Patrick C. Walsh Prostate Cancer Research Fund and a Prostate Cancer Foundation Young Investigator Award to K.S.S. We thank the Yegnasubramanian Lab for providing access to Droplet Digital equipment and Dr. Michael Haffner for helpful discussion. The authors acknowledge the Sidney Kimmel Comprehensive Cancer Center (SKCCC) Experimental Irradiator Core for providing access to the CIXD Biological Irradiator and thank Esteban Velarde for technical assistance.

