## Supplemental Figures S1-S2; Supplemental Tables S1-S3 for "Use of Droplet Digital PCR for Consistent Detection of TMPRSS2:ERG Gene Fusion Transcripts Initiated In Vitro"

### Supplemental Figures and Supplemental Figure Legends

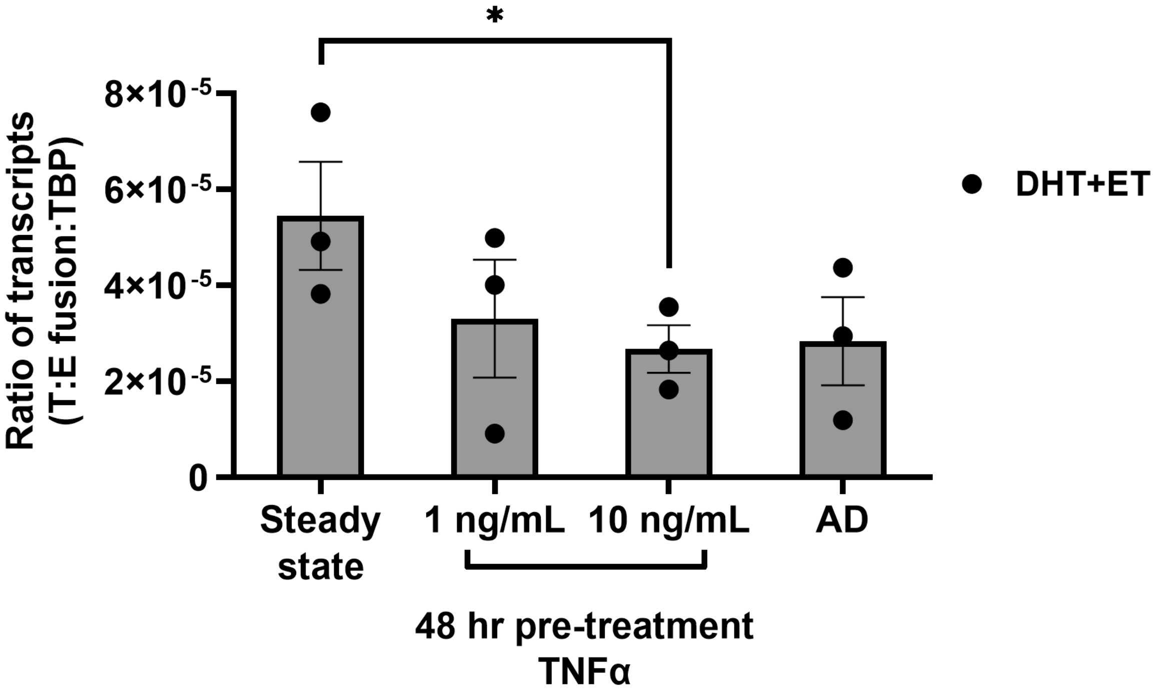


**Supplemental Figure S1.** LNCaP cells pre-treated with TNFα and then exposed to DHT and etoposide (DHT+ET) show a reduced ratio of T:E fusion transcripts relative to TBP reference gene transcripts. LNCaP cells were pre-treated with TNFα (1 or 10 ng/mL), androgen deprivation (AD, RPMI + 10% CSS), or steady state media (RPMI + 10% FBS) for 48 hours prior to exposure to DHT (100 nM) and etoposide (100 µM) for T:E gene fusion generation. The concentration of T:E gene fusion transcripts was determined by ddPCR. Ratio was calculated by dividing the concentration (copies/µL) of T:E gene fusion transcripts by TBP transcripts. *TBP* served as a reference gene. Data is representative of 3 biological replicates. Ratio is displayed as mean +/- SEM. Statistical significance was assessed using a one-tailed unpaired t test, with significance set at $p < 0.05$. ($^*p < 0.05$).

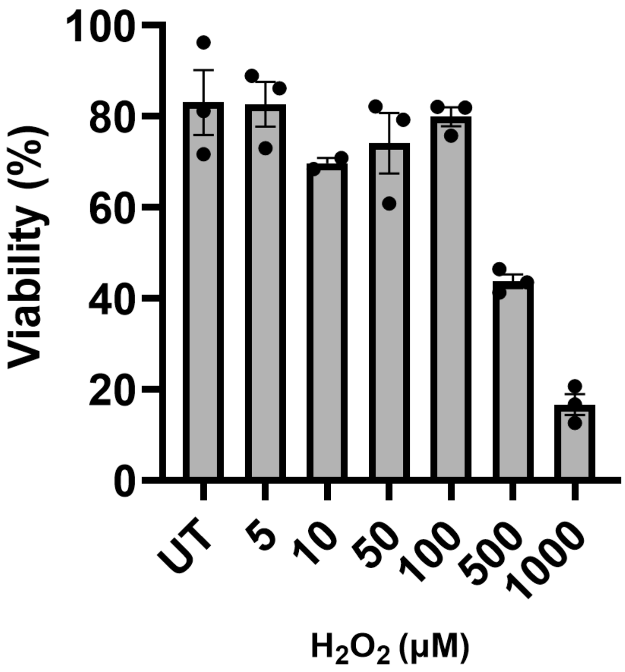


**Supplemental Figure S2.** LNCaP cell viability after exposure to $H_2O_2$. LNCaP cells were exposed to $H_2O_2$ (5-1000 μM) for 24 hours. Upon harvest, cells were diluted 1:1 with trypan blue and counted using the Countess Automated Cell Counter. Two technical replicates were performed for cell counting. Viability was determined by dividing the average number of live cells (trypan blue-) by the averaged total number of cells (trypan blue+/-) and multiplying by 100. Data is representative of 3 biological replicates. Viability is displayed as mean +/- SEM. UT = untreated control.

**Supplemental Table S1.** Cycle conditions for Droplet Digital PCR (ddPCR).

| | Temperature (°C) | Time | Number of cycles |
|---|---|---|---|
| Enzyme activation | 95 | 10 min | 1 |
| Denaturation | 95 | 30 s | 39 |
| Annealing/extension | 60 | 1 min | |
| Enzyme deactivation | 98 | 10 min | 1 |
| End/Hold | 12 | ∞ | 1 |

**Supplemental Table S2**. Cycle conditions for reverse transcription quantitative PCR (RT-qPCR).

| | Temperature (°C) | Time | Number of cycles |
|---|---|---|---|
| Enzyme activation | 95 | 30 s | 1 |
| Denaturation | 95 | 3 s | 40 |
| Annealing/extension + plate read | 60 | 22 s | |

**Supplemental Table S3**. Sequence of TMPRSS2:ERG gBlocks® gene fragment (IDT).

| TMPRSS2:ERG gene fragment |
|---|
| 5'—<br>ATATGATCGATCGAGTAGGCGCGAGCTAAGCAGGAGGCGGAGGCGGAGGCGGAGGGCGAGGG<br>GCGGGGAGCGCCGCCTGGAGCGCGGCAGGAAGCCTTATCAGTTGTGAGTGAGGACCAGTCGTT<br>GTTTGAGTGTGCCTACGGAACGCCACACCTGGCTAAGACAGAGATGACCGCGTCCTCCTCCAGC<br>GACTATGGACAGACTTCCAAGATGAGCCCACGCGTCCCTCAGCAGGATTGGCTGTCTCAACCCC<br>CAGCCAGGGTCACCATCAAAATGGAATGTAACCCTAGCCAGGTGAATGGCTCAAGGAACTCTCC<br>TGATGAATGCAGTGTGGCCAAAGGCGGGAAGATGGTGGGCAGCCCAGACACCGTTGGGATGAA<br>CTACGGCAGCTACATGGAGGAGAAGCACATGCCACCCCCAAACATGACCACGAACGAGCGCAG<br>AGTTATCGTGCCAGCAGATCCTACGCTATGGAGTACAGACCATGTGCGGCAGTGGCTGGAGTGG<br>GCGGTGAAAGAATATGGCCTTCCAGACGTCAACATCTTGTTATTCCAGAACATCGATGGGAAGG<br>A—3' |
